# Left angular gyrus disconnection impairs multiplication fact retrieval

**DOI:** 10.1101/2021.10.27.465904

**Authors:** S. Smaczny, C. Sperber, S. Jung, K. Moeller, H.-O. Karnath, E. Klein

## Abstract

Arithmetic fact retrieval has been suggested to recruit a left-lateralized network comprising perisylvian language areas, parietal areas such as the angular gyrus (AG), and subcortical structures such as the hippocampus. However, the underlying white matter connectivity of these areas has not been evaluated systematically so far.

Using simple multiplication problems, we evaluated how disconnections in parietal brain areas affected arithmetic fact retrieval following stroke. We derived disconnectivity measures by jointly considering data from n=73 patients with acute unilateral lesions in either hemisphere and a white-matter tractography atlas (HCP-842) using the Lesion Quantification Toolbox (LQT). Whole-brain voxel-based analysis indicated a left-hemispheric cluster of white matter fibers connecting the AG and superior temporal areas to be associated with a fact retrieval deficit. Subsequent analyses of direct grey-to-grey matter disconnections revealed that disconnections of additional left-hemispheric areas (e.g., between the superior temporal gyrus and parietal areas) were significantly associated with the observed fact retrieval deficit.

Results imply that disconnections of parietal areas (i.e., the AG) with language-related areas (i.e., superior and middle temporal gyri) seem specifically detrimental to arithmetic fact retrieval. This suggests that arithmetic fact retrieval recruits a widespread left-hemispheric network and emphasizes the relevance of white matter connectivity for number processing.

## 1. Introduction

The dominating view in numerical cognition is that arithmetic facts such as multiplication tables are stored in and retrieved from long-term memory in a verbal format (e.g., Dehaene et al., 2003; Delazer et al., 2003; Grabner et al., 2009). The triple code model (Dehaene & Cohen, 1995; 1997; Dehaene et al., 2003), which has been remarkably successful in providing a theoretical framework for neuroimaging research on numerical cognition, posits that the retrieval of arithmetic facts recruits areas also involved in language processing such as left-hemispheric perisylvian areas and the left angular gyrus (AG). In contrast, the manipulation of numerical content during magnitude-manipulation based calculation (e.g., addition, subtraction) has been associated with a fronto-parietal network for number (magnitude) processing centered around the intraparietal sulcus (IPS; e.g., Dehaene et al., 2003; for a meta-analysis see Arsalidou & Taylor, 2011). This network has recently been coined the ‘math-responsive network’ (Amalric & Dehaene, 2017; 2019).

The retrieval of rote learned arithmetic facts has been assumed to be processed multi-modularly and distributed within a left-lateralized language network, including inferior frontal gyrus (IFG; Delazer et al., 2003), middle and superior temporal gyri (MTG/STG), supramarginal gyrus (SMG), angular gyrus (AG; e.g., Dehaene et al., 2003; Delazer et al., 2003; Grabner et al., 2009a; 2009b), and hippocampus (e.g., Bloechle et al., 2016; Klein et al., 2016; 2019). However, the exact role of some of these areas is still debated. On the one hand, several case studies reported patients with a preserved AG who presented selective deficits for arithmetic fact retrieval (as measured by a multiplication task, Cohen et al., 2000; Dehaene & Cohen, 1997; Van Harskamp et al., 2005). Moreover, the role of the hippocampus, which is assumed a key structure for long-term memory in general, is not yet fully understood.

Delazer and colleagues (2019), for instance, suggested that the hippocampus plays a crucial role in encoding and consolidating arithmetic facts, but not in their actual retrieval after consolidation in memory. On the other hand, it has been criticized that the interaction of several of these areas, which are activated during arithmetic fact retrieval, may not be specific to arithmetic fact retrieval. For instance, IFG, STG, and SMG are typically also involved in phonological decoding during language processing (Prado et al., 2011; Vigneau et al., 2006; for a review, see Price, 2012). Similarly, modulation of AG (de)activation, observed for arithmetic fact retrieval, was also found for non-mathematical content (Ischebeck et al., 2007; Zamarian et al., 2009). Therefore, it appears likely that (sub)processes of arithmetic fact retrieval are related to language processing.

However, such an association between arithmetic fact retrieval and language processing is still debated. Amalric and Dehaene (2016, 2017, 2019) suggested that simple, overlearned arithmetic facts might be processed independently of left-hemispheric language areas. In particular, the authors proposed that procedure-based calculation and overlearned arithmetic facts are processed within the bilateral fronto-parietal ‘math-responsive’ network.

However, the empirical evidence concerning this question is still inconsistent on whether arithmetic fact retrieval and language processes recruit overlapping cortical circuits. There seems to be a consensus that arithmetic fact retrieval is processed multi-modularly and distributed in distinct brain regions. Therefore, it is critical to consider network characteristics such as connectivity between distant brain regions and subnetworks.

Klein et al. (2016) provided first evidence for white matter connectivity underlying arithmetic fact retrieval using probabilistic fiber tracking in healthy participants. The authors showed that parietal areas are connected with frontal areas dorsally via the cingulate bundle and ventrally via the extreme/external capsule system and temporal areas via the inferior longitudinal fascicle. The importance of the integrity of such a network architecture for fact retrieval has already been highlighted in some studies: For instance, in a re-evaluation of a single case reported by Zaunmueller et al. (2009), Klein et al. (2013b) found that the lesion was limited to the basal ganglia region (and did not involve the AG) but nevertheless led to disruption of white matter fibers connecting frontal areas with the AG. Based on this, the authors suggested that the patient could not retrieve arithmetic facts because of a disconnection within their retrieval network and not because of actual grey matter damage in cortex areas associated with fact retrieval. In this vein, Mihulowicz et al. (2014) conducted a voxel-based lesion behaviour mapping study (VLBM). They observed a significant association of lesions in the dorsal pathway (i.e., parts of the superior lateral fasciculus) with arithmetic fact retrieval.

Only recently, lesion-disconnectome mapping has brought forward the possibility to more closely examine the role of network disruption of brain lesions in post-stroke cognitive deficits. In particular, it allows for an indirect assessment of lesion-induced disconnections by reference to healthy connectome data (e.g., Griffis et al., 2019). In the current study, we evaluated brain disconnections associated with a deficit in arithmetic fact retrieval in acute left- and right-hemisphere stroke patients at the group level. We expected that white matter disconnections within the left-hemispheric arithmetic fact retrieval network, including, among others, AG, MTG, and hippocampus, should be associated with deficits in arithmetic fact retrieval. In a subsequent analysis, we further aimed at evaluating which disconnections between two specific grey matter regions are significantly associated with a deficit in arithmetic fact retrieval.

## 2. Methods

### 2.1 Participants

We used the existing patient data set from Mihulowicz et al. (2014) and added 28 new patients for the present analyses. In total, 73 native German-speaking acute stroke patients participated in the study. All of them were patients consecutively admitted to the Center of Neurology at Tübingen University Hospital over 33 months (cf. Mihulowicz et al., 2014) plus another 18 months. Among these, 38 patients presented left hemisphere damage (LHD) and 35 patients right hemisphere damage (RHD). Patients were only included when they presented an MR- or CT-documented cerebral stroke, not more than 14 days post-stroke. We did not include patients with previous lesions, other neurological or psychiatric diseases, substantial micro-angiopathy or white matter alterations. Patients or their relatives gave their informed consent to participate in the study. The study followed the ethical standards in the Declaration of Helsinki (Version 2013) and was approved by the local ethics committee (Vote 82/2018 BO2). Clinical and demographic data for the whole group of 73 patients are given in Table 1; an illustration of lesion overlap of all participants can be seen in Figure 1.

**Table 1.** Demographic and clinical data of all LHD and RHD patients.

|  |  | **LHD** | **RHD** |
| --- | --- | --- | --- |
| *N* |  | 38 | 35 |
| Sex |  | 19F, 19M | 18F, 17M |
| Age (years) | Mean (SD) | 63.05<br>(14.22) | 63.38<br>(15.37) |
| Stroke type | Ischemic / Hemorrhage | 32 / 6 | 28 / 9 |
| Interval lesion onset to examination (days) | Mean (SD) | 4.8 (2.5) | 4.4 (2.3) |
| Interval lesion onset to imaging (days) | Mean (SD) | 2.7 (2.6) | 2.7 (3.2) |
| Education (years) | Mean (SD) | 11.27 (4.36) | 11.14 (4.43) |
| Contralateral paresis | % present | 52.63 | 65.71 |
| Visual field deficit | % present | 13.16 | 11.43 |
| Aphasia | % present | 42.10 | - |
| Spatial Neglect | % present | - | 13.89 |

**FIGURE 1.**
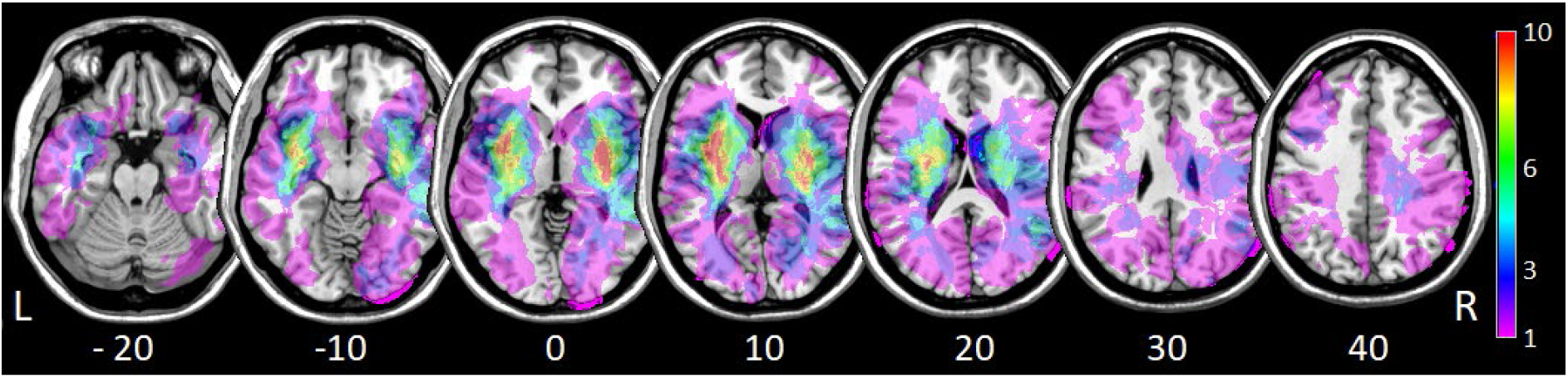
Lesion Overlap. Overlap of all lesions included in the analysis. The color bar indicates the frequency of lesion overlap. The vertical z coordinate of standardized MNI space is given below each slice.

### 2.2 Procedure

Patients were investigated in sitting position. The entire experiment took about one hour and was completed in one session for most patients. For some, further sessions were needed due to clinical examinations or because they needed a break. For the multiplication task, patients responded verbally and in written form when the former was not possible.

We ensured that all patients were able to follow the task instructions. In particular, we tested receptive and expressive language abilities in LHD patients. For *language comprehension*, we used the ‘Colour-Figure subtest from the German adaptation of the Aphasia Check List (ACL; Kalbe et al., 2005). For *language production*, we applied the ‘Picture naming task’ of the Aachener Aphasia-Bedside Test (AABT; Biniek et al., 1992).

We also tested patients with RHD for *spatial neglect*. The neglect test comprised the ‘Letter Cancellation Task’ (Weintraub & Mesulam, 1985), the ‘Bells test’ (Gauthier et al., 1989), as well as a copying task (Johannsen & Karnath, 2004). Patients were diagnosed with spatial neglect if they scored above the respective cut off as per manual in at least 2 out of 3 tests. We calculated the Centre of Cancellation (CoC) for the two cancellation tasks, using the procedure and cut-off scores for diagnosing spatial neglect described by Rorden and Karnath (2010).

In the copying task, patients were asked to copy a complex multi-object scene consisting of four figures (a fence, a car, a house, and a tree), with two of them located in each half of the horizontally oriented sheet of paper. Omission of at least one of the contralateral features of each figure was scored as 1. The omission of each whole figure was scored as 2. One additional point was given when contralaterally located figures were drawn on the ipsilesional side of the paper sheet. The maximum score was 8. A score higher than 1 (i.e. > 12.5% omissions) was taken to indicate neglect.

Visual field deficits were assessed in all patients via the standard confrontational procedure. Table 1 presents all clinical data.

### 2.3 Stimuli

As part of our battery, we evaluated patients’ arithmetic fact retrieval through a single-digit multiplication production task (i.e., 42 items from arithmetical tables up to 9×9), included in the standardized neuropsychological Number Processing and Calculation Battery (NPC; Delazer et al., 2003). Each multiplication problem was presented on a separate A4 sheet (black digits printed on white paper, digit height: 7 mm). Sheets were aligned centrally on a table in front of the patient. To operationally determine impaired performance, we used the cut-off criteria provided for the NPC (cf. Delazer et al., 2003). According to the standardized NPC procedure, testing was stopped after five consecutive incorrect or missing responses, and no time limit was imposed. However, responses lasting longer than 10 seconds were rated as incorrect, as this implies a failure in instant fact retrieval. Self-corrections were allowed.

One aphasic LHD patient could not solve the single-digit multiplication production task orally or in writing, while he could solve 24 of 25 addition tasks correctly in verbal form. In this patient, we applied a multiple-choice version of the multiplication task also provided in the NPC to determine a core deficit in multiplication processing rather than a language difficulty. The multiple-choice version tested the same multiplication problems (e.g., 6 × 7) by providing four solution probes (i.e. correct solution, 42; one operand error, e.g., 45, one table error, e.g., 49, and one close-miss error, e.g., 39). The patient showed deficits in the multiple-choice version. Accordingly, he was included in data analysis.

### 2.4 Lesion Analysis

We used diffusion-weighted MRI images taken within 48h after stroke (Weber et al., 2000) or else T2-weighted fluid-attenuated inverse-recovery (FLAIR) images (Brant-Zawadzki et al., 1996; Noguchi et al., 1997; Ricci et al., 1999; Schaefer et al., 2002). In case MR imaging was not conducted, CT images were used (MRI: *N*=29; CT: *N*=44). When several imaging data sets existed for an individual patient, we used the session closest in time to behavioral testing showing a clear demarcation. Lesion borders were marked semi-automatically using the ‘Clusterize Toolbox’ (de Haan et al., 2015). Then, both the anatomical scan and the lesion map were normalized into stereotaxic space using the ‘Clinical Toolbox’ (Rorden et al., 2012; www.mccauslandcenter.sc.edu/CRNL/clinical-toolbox) implemented in SPM12 (www.fil.ion.ucl.ac.uk/spm).

### 2.5 Whole-brain disconnectivity mapping

We created individual white matter disconnectivity topographies for each patient. These topographies indicate the proportion of disconnected fibers for each white matter voxel in the brain imaging space running through this voxel. Thereby, it allows a topographical assessment of a lesion’s impact on brain connectivity as a whole. To this end, we applied the Lesion Quantification Toolkit (LQT; Griffis et al., 2021). The LQT utilizes a tract-wise connectome atlas and embeds the patient’s lesion map into it. The toolkit identifies all fiber streamlines that intersect with the lesion and maps connectome-wide disconnection induced by the lesion. We used the LQT’s standard HCP-842 atlas (Yeh et al., 2018) for atlas-based tractography, keeping the default parameter setting as suggested in the LQT manual.

Subsequently, we further analyzed these continuous disconnectivity maps using mass-univariate General Linear Models in ‘NiiStat’ (https://github.com/neurolabusc/NiiStat). Only voxels with a disconnection in at least five patients were considered in the analysis. Tests were performed one-sided at *p*<0.05 and corrected for family-wise errors. The family-wise error rate was corrected using 5000 permutations with maximum statistic permutation (Nichols & Holmes, 2002). This disconnectivity topography mapping thus identified voxels for which disconnection is associated with a more severe deficit in multiplication.

### 2.6 Region-to-region disconnectivity

We additionally analyzed parcel-wise disconnectivity as provided by the LQT. This procedure allowed us to identify which direct disconnections between two grey matter regions are significantly associated with a deficit in the multiplication task. We created a structural connectivity matrix by combining the provided tractography atlas and the Brainnetome Atlas (BN-246, Fan et al., 2016) as our parcellation atlas. The BN-246 is multi-modally derived, contains 210 cortical and 36 subcortical subregions and was developed specifically for connectivity analyses. The number of streamlines disconnected by the lesion map between each pair of parcels was converted to a percentage of disconnected streamlines, resulting in symmetric 246-by-246 disconnectivity matrices. We subjected these matrices to a mass-univariate analysis to identify associations between disconnection and the deficit using custom scripting in MATLAB R2020a. Following the strategy of the topographical analysis of brain-wide disconnectivity maps described above, we employed mass-univariate general linear models with a family-wise error correction by maximum statistic permutation (Nichols & Holmes, 2002).

We loaded disconnection matrices into MATLAB and removed the diagonal and redundant elements below it. Many ROI-to-ROI disconnections were rarely or never present in the data, likely either because the sample’s lesion anatomy did not include damage to the connection or because the connection was physiologically non-existent. Therefore, we identified the sum of all patients with a disconnection present (i.e., a disconnection score > 0) for each ROI-to-ROI connection. We removed all connections affected in less than 15 patients from the analysis, reducing the final set of analyzed connections to 955. We then computed a general linear model for each ROI-to-ROI connection with the independent variable disconnectivity score and the dependent variable multiplication performance. Then, maximum statistic permutation with 50.000 permutations was employed on permuted behavioral data and the original disconnection data with the same analysis strategy to assess the distribution of maximum statistics under the null hypothesis. We obtained a one-sided, corrected threshold for statistical significance at *p* < 0.05 by identifying the 95^th^ percentile of permutation-derived maximum statistics. To identify those connections most strongly associated with the deficit, we then identified a smaller set of the strongest disconnection-deficit associations at *p* < 0.01.

### 2.7 Data availability

Online materials, including all analysis scripts, descriptive data, and resulting topographies, are publicly available at http://dx.doi.org/10.17632/yjkr647mzb.1. The clinical datasets analyzed in the current study are not publicly available due to the data protection agreement approved by the local ethics committee and signed by the participants.

## 3. Results

### 3.1 Behavioral measures

Table 2 summarizes the behavioral results of the multiplication task. Items involving ‘0′ or ‘1 as operands (n=6) were excluded from the analysis because they address rule-based processing (McCloskey et al., 1991). While the analysis used continuous values, five patients in the RHD group performed below the typical cut-off in the multiplication task; in the LHD group, ten patients did so, including one aphasic patient who could not respond to the multiplication task orally or in writing. They were presented with a multiple-choice version (see above).

**Table 2.**
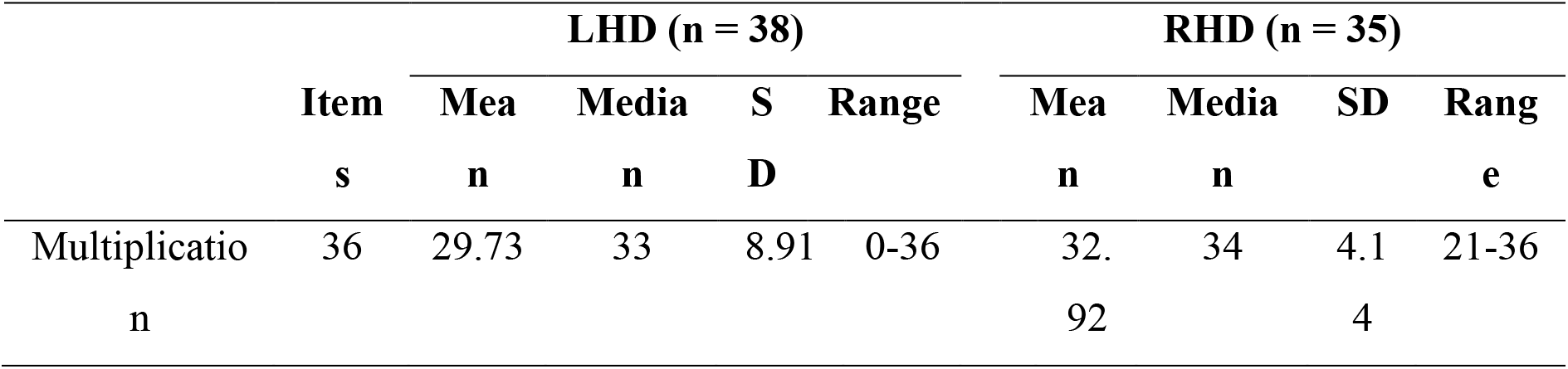
Raw scores (number of items solved correctly) observed for the two patient groups in the multiplication task.

### 3.2 Brain-wide disconnectivity mapping

Figure 2 (Panel A) illustrates the average percentage of lesion-induced disconnection for all 73 patients. Deficits in single-digit multiplication were significantly associated with the disconnection of a single cluster around × = (−59; −40), y = (−57; −39), and z = (3; 22), affecting areas defined by the HCP-842 as the left arcuate fasciculus, temporopontine tract, and U-fibers between the AG and superior temporal areas (see Figure2 (Panel B).

**FIGURE 2.**
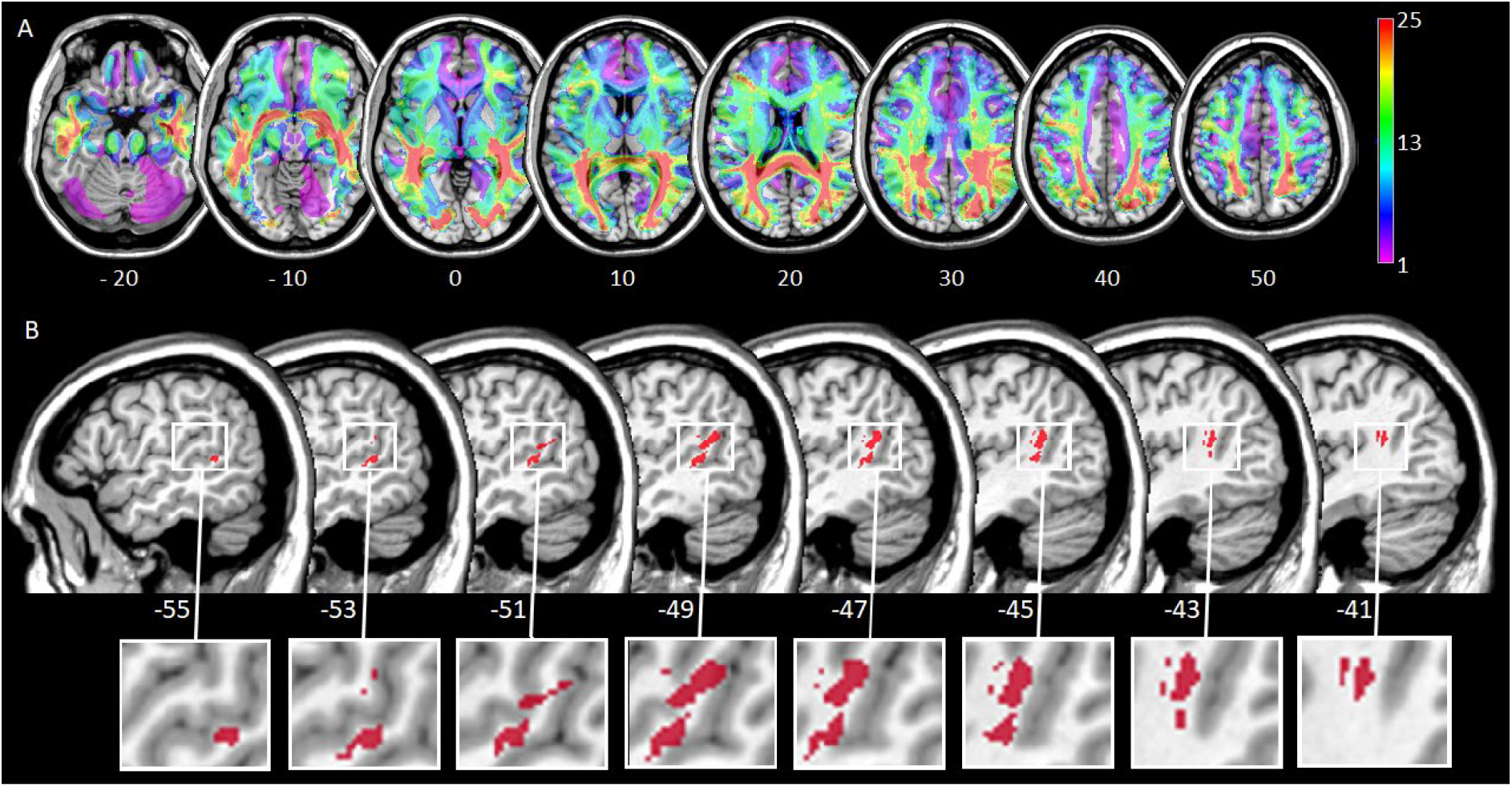
Disconnectivity mapping. (A) The percentage of fibre disconnection as indicated by the average reduction in streamline density is color-coded from 1 (pink) to 25 (red), whereby the maximum value in a patient was 36. A higher number indicates a higher percentage of fibre disconnection. Topographies can be viewed in the online data. The vertical z coordinate of standardized MNI space is given below each slice. (B) Sagittal view of the left hemisphere. Statistical voxel-wise lesion-behavior mapping (VLBM) analyses using mass-univariate general linear models in multiplication. Plotted are voxels that survived permutation-test based FWE correction at *p* < 0.05 (left hemisphere). The sagittal × coordinate of standardized MNI space is given below each slice.

### 3.3 Region-to-region disconnectivity

Mapping multiplication score to ROI-to-ROI disconnectivity using general linear models identified 36 connections significant at *p* < 0.05 (see Figure 3 (Panel A), Table 3). The disconnectome revealed several regions that stood out due to several disconnections associated with the multiplication deficit. A hub-like structure was found for the left thalamus, with 17 disconnections mainly to the left inferior and superior parietal lobules, including PGa of the angular gyrus. Furthermore, 6 disconnections between the left IFG and the medioventral occipital cortex as well as the left lateral occipital cortex were also associated significantly with poorer multiplication scores. Finally, further 5 disconnections from the left STG to the right inferior and superior parietal lobules (IPL, SPL), as well as to the right precuneus, were significant.

**Table 3.**
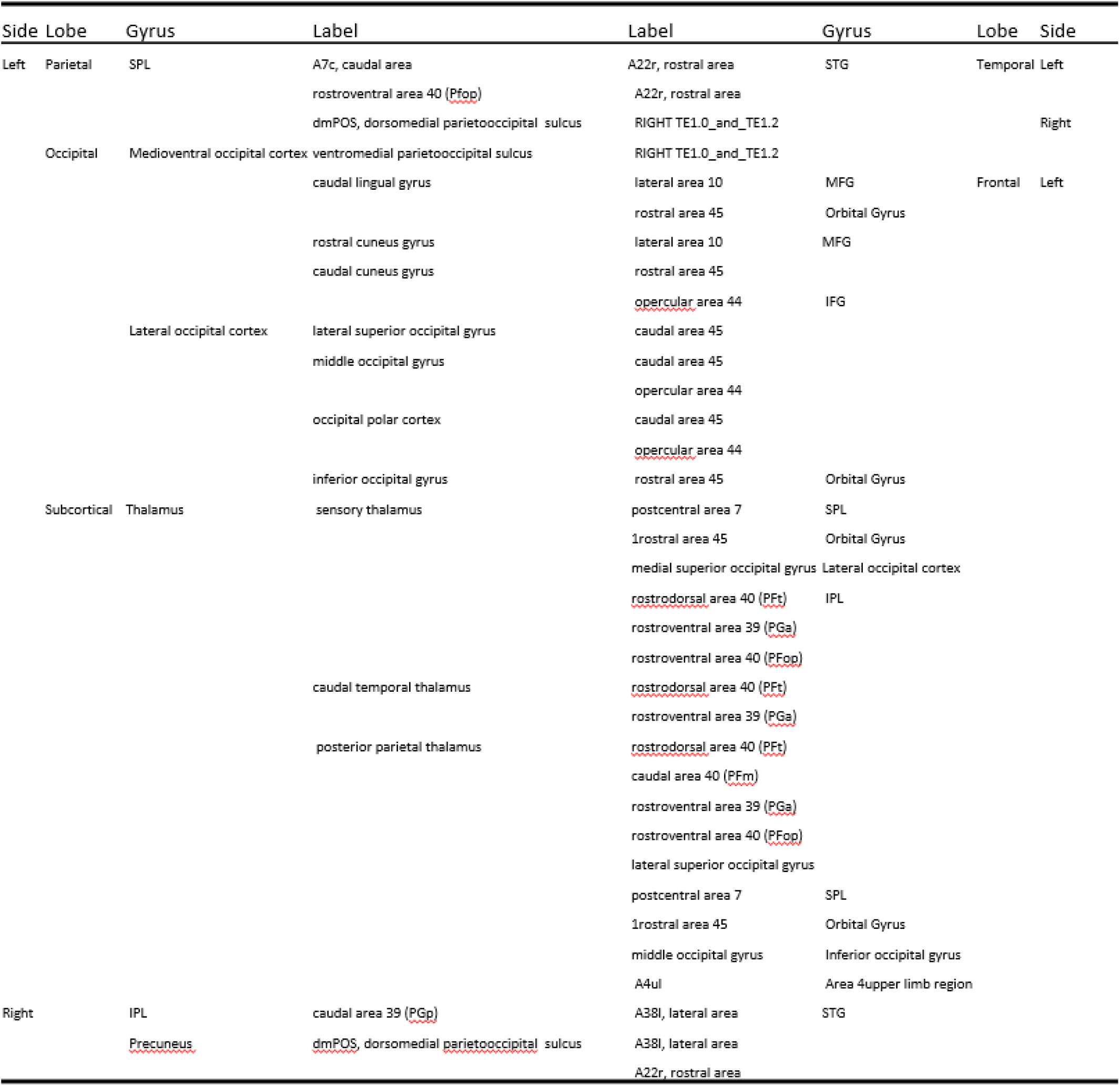
Parcel-to-parcel disconnections following the region-to-region disconnectivity analysis at *p* < .05.

**Figure 3.**
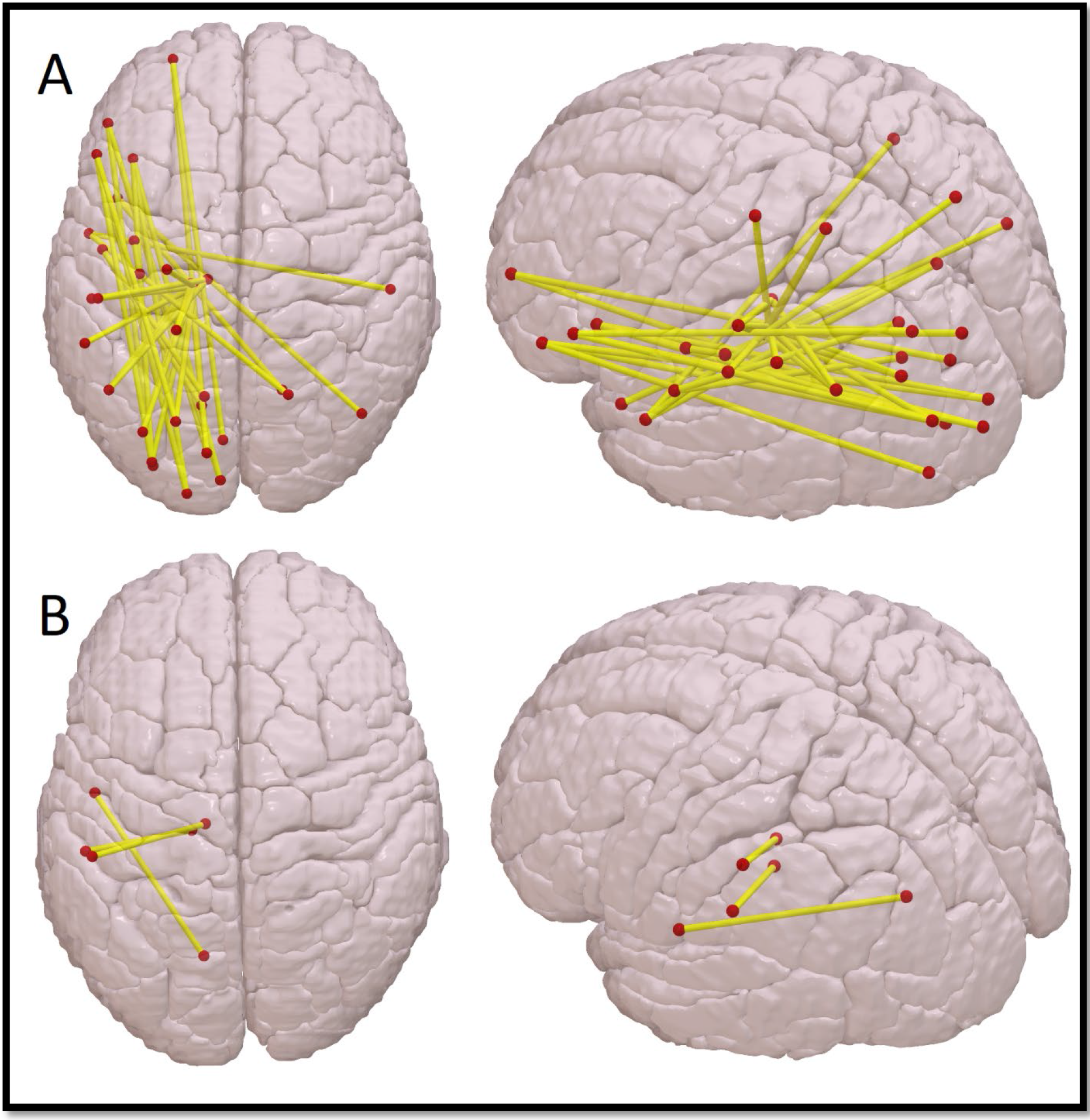
Parcel-to-parcel disconnections following the region-to-region disconnectivity analysis. (A) presents the disconnections significant at *p* < .05, (B) those at *p* < .01.

To facilitate the interpretation of the strongest predictors, we then set the threshold to *p* < 0.01. This procedure revealed a small set of 3 left-hemispheric disconnections to have the strongest association with poorer multiplication scores (see Fig. 3 (Panel B)): Te1.0 / Te1.2 of the primary auditory cortex with the ventromedial parietooccipital sulcus, posterior parietal thalamus with IPS (rostroventral area 40) and the left caudal temporal thalamus with the IPS (rostrodorsal area 40).

## 4. Discussion

The present study evaluated how white matter disconnection affected arithmetic fact retrieval. We carried out two lesion-disconnection analyses in first-time stroke patients with unilateral damage to either the left or right hemisphere performing a multiplication task. In the first analysis, patients’ lesions were superimposed to a white matter tractography map, leading to individual disconnection topographies (Griffis et al., 2019). A whole-brain univariate analysis suggested that disconnection of white matter fibers situated between the anterior AG and posterior STS was associated with a multiplication deficit. A second analysis was carried out to investigate direct disconnections between two brain areas implicated in multiplication deficits. Single brain areas were defined as ROIs by a connectivity-based grey matter atlas, and disconnection between each pair of regions was investigated for its role in multiplication deficits. We found several structures with multiple relevant disconnections. These included the left thalamus, (intra)parietal areas such as the AG, STG, IFG, and occipital areas. Single disconnections with the strongest disconnection-deficit association were found between the thalamus and the IPS as well as occipital regions, but also between the primary auditory cortex and occipital areas. Results of these analyses will be discussed in turn in the following.

### Whole-Brain Analysis: Disconnection of the AG with temporal areas

The findings of the whole-brain analysis fit well with a connectivity approach of arithmetic fact retrieval put forward by Klein et al. (2016). In this approach, connections between the MTG and the AG were considered necessary for arithmetic fact retrieval, as both areas are essential elements of a left-hemispheric arithmetic fact retrieval network. A significant role of the AG for arithmetic fact retrieval has also been suggested by several neuropsychological single case studies on arithmetic fact retrieval (e.g., Hittmair-Delazer et al., 1994; Lee, 2000; Cohen et al., 2000). However, other single case studies have questioned the importance of the AG within this network, as they reported patients presenting with a multiplication deficit despite having a preserved AG (Cohen et al., 2000; Dehaene & Cohen, 1997; van Harskamp et al., 2005; Zaunmueller et al., 2009). For example, patient ATH had severe difficulties verbally solving multiplication problems while making only a few mistakes in subtraction (Cohen et al., 2000). On the other hand, Van Harskamp & Cipolotto (2001) reported on a patient who did not show a multiplication deficit, although damage to left parietal regions, including the AG, occurred.

Given both the current results and more recent research, inconsistent results of single case studies can be explained by the approach of Klein and colleagues (2016): The disconnection cluster indicated by our analysis to be associated with arithmetic fact retrieval involved the left arcuate fasciculus, the temporopontine tract and U-fibers between the AG and STS. This observation suggests that disconnection of the AG with temporal areas, such as STS, STG, and MTG seems to cause an arithmetic fact retrieval deficit. This expands the results of previous studies indicating the detrimental effect of lesions to or disconnection of the AG. In this sense, Klein et al. (2013b) re-evaluated a single case reported by Zaunmueller et al. (2009), focusing on lesions to the patient’s white matter compared to previous single case studies focusing on lesions more on grey matter damage. While the lesion itself was in the basal ganglia, it also incorporated white matter fibers that explicitly connect frontal areas with the AG, suggesting that the patient could not retrieve arithmetic facts due to poor connectivity of the AG. Similarly, Mihulowicz et al. (2014) conducted a voxel-based lesion behavior mapping study observing that white matter lesions to superior parts of the superior lateral fasciculus (SLF II) impaired arithmetic fact retrieval.

The current findings provide first evidence on explicit measures of disconnection on arithmetic fact retrieval on the group level, further corroborating the role of the AG within the arithmetic fact retrieval network. They can explain case studies questioning the role of the AG in multiplication, whereby patients with multiplication deficits without damage to the AG or other grey matter areas of the network may have suffered from structural white matter disconnectivity within the network, as suggested by Klein et al. (2013b, c).

Importantly, neurofunctional evidence from healthy participants on learning complex multiplication facts (e.g., Delazer et al., 2003, 2005; Ischebeck et al., 2006, 2007; Grabner et al., 2009; Rickard et al., 2000; Zamarian et al., 2009) also indicates that the AG plays a crucial role in arithmetic fact retrieval. Bloechle et al. (2016) evaluated fMRI signal before and after multiplication training. They found a stronger fMRI signal in the AG when comparing trained vs. untrained multiplication tasks after training only, but no difference in activation when directly comparing the same multiplication problems across training (post- vs. pre-training). Therefore, the authors suggested that the AG might act as a ‘circuit breaker’, coordinating whether a task is to be solved purely via fact retrieval or whether a switch in strategy requiring additional cognitive processing, such as calculation, is necessary.

This aspect provides a compelling explanation for patient FS’s ability to multiply correctly despite damage to parietal areas, including the AG: They were still able to recall multiplication facts due to an intact temporal retrieval network and did not require the AG to switch from fact retrieval to calculation. In the current study, most patients with an arithmetic fact retrieval deficit also suffered from aphasia. Accordingly, they might have not been able to recall phonologically stored arithmetic facts initially. When attempting to switch to a calculation-based strategy, these patients were also slowed due to the disconnection of AG. Therefore, a possible explanation for the given findings may be that when arithmetic fact retrieval fails, the AG is involved in switching the strategy by which a task is solved. When it is disconnected or damaged, this switch fails.

### Region-to-Region Disconnectivity

Results of the second analysis indicate several disconnections between left-hemispheric areas, which lead to deficits in arithmetic fact retrieval. When considering the analysis using the more liberal threshold (Figure 3, Panel A), several disconnection-groupings were significant. First off, there were several disconnections of the left thalamus with parietal areas. The thalamus is involved in various cognitive processes, such as memory, language or mental set-shifting (see Saalmann & Kastner; 2015 for an overview). It is also explicitly considered to be involved in mental arithmetic (Arsalidou & Taylor, 2011; Johnson & Ojemann, 2000; Koyama et al., 2020). In the more conservative threshold analysis, two of the three significant disconnections were of the thalamus involved connections with the intraparietal sulcus (IPS).

There is a broad consensus that the IPS is involved in magnitude processing (for meta-analyses see Arsalidou & Taylor, 2011; Arsalidou et al., 2018; Dehaene et al., 2003; Hawes et al., 2019). Yet, arithmetic tasks are typically not solved exclusively by either arithmetic fact retrieval or magnitude manipulations (i.e., calculations). Instead, they are solved by an adaptive interplay between both (Klein et al., 2016). It thus seems possible that arithmetic fact retrieval deficits following disconnection between the thalamus and the IPS reflect difficulties in switching between and integrating fact retrieval and magnitude manipulation strategies as already argued on the whole-brain level. As a general conclusion, the thalamus may be considered as a sort of central relay of information between different cortical areas, receiving, coordinating and transmitting information across the cortex. Therefore, it should not be surprising that disconnection of such a vital structure with anatomical correlates associated with number processing, such as left-hemispheric parietal areas, would lead to deficits in multiplication.

Our results showed several disconnections of the left IFG with left (parieto-)occipital areas. While previous research reported essential connections between parietal regions, such as the AG, and the IFG, the role of fronto-occipital connections in arithmetic has not been focussed on so far. Yet, it is known that the IFG is connected with the parietal and occipital cortex via the inferior fronto-occipital fasciculus (IFOF). This suggests that disconnection of certain parts of the IFOF may also lead to arithmetic fact retrieval deficits. The EC/EmC system corresponds to a rostral/anterior segment of the IFOF (Willmes et al., 2014) and is considered to be part of a ventral pathway involved in arithmetic fact retrieval (Klein et al., 2016). More precisely, Klein et al. (2013b) found that arithmetic tasks which can be solved via arithmetic fact retrieval were associated with a ventral processing route. This route ranges from frontal regions, such as the medial frontal gyrus to parietal regions, including the AG via the middle longitudinal fascicle (MdLF), which itself is part of the EC/EmC system. They suggested that this connection serves the purpose of phonological or semantic access, similar to language processes (e.g., Saur et al., 2008; Weiller et al., 2011). Therefore, in the current study, the observed disconnections might have led to language-related errors, such as semantic or phonemic paraphasias.

Finally, we observed several disconnections of the STG associated with an arithmetic fact retrieval deficit. Most of these disconnections were to the SPL, more specifically, the precuneus. This observation fits well with a series of fMRI studies in healthy adults (Prado et al., 2011) as well as children with and without mathematical learning disabilities (e.g., Berteletti et al., 2011; Demir et al., 2014) that suggest that the STG, as well as the MTG play a role in arithmetic fact retrieval. Similarly, the more conservative threshold analysis also found a disconnection between the primary auditory cortex and the ventromedial parietooccipital sulcus to be associated with impaired arithmetic fact retrieval. These connections have been suggested to run via the MdLF, emphasizing its importance in the arithmetic fact retrieval network.

### The role of the AG in arithmetic fact retrieval

Taken together, the current results indicate that arithmetic fact retrieval is subserved by a widespread left-hemispheric network including the AG, superior temporal gyrus, and the thalamus, among others, thus corroborating the network suggested by Klein et al. (2016). In particular, the first analysis corroborated the idea that the angular gyrus plays a central role in arithmetic fact retrieval (e.g., Delazer et al., 2003; Dehaene et al., 2003) as a disconnection of the AG led to impaired arithmetic fact retrieval. However, the second analysis revealed that not only AG connectivity but also connectivity of various other left-hemispheric areas, including the IPS, seems to be necessary for the successful retrieval of arithmetic facts. As regards the role of the AG in the fact retrieval network, we suggest that the AG may not be the (sole) storage location of arithmetic facts. Instead, it may serve a more regulatory role within the network, for instance, subserving semantic integration of concepts (Amalric & Dehaene, 2017; Price et al., 2015) or strategy switching (Bloechle et al., 2016). For instance, patients who could not solve tasks via arithmetic fact retrieval may have had to switch to a different strategy requiring increased magnitude processing (e.g., Dehaene et al., 2003; Klein et al., 2016). If these processing routes to the IPS of either hemisphere were also disconnected (i.e., due to a lesion affecting both association and commissural fibers), patients might not have been able to use this compensatory mechanism adequately. Consequentially, they committed errors or took longer than the ten-second cut-off time to solve the task.

## Conclusion

Our data on lesion-disconnection analyses in unilateral stroke patients suggest that a deficit in arithmetic fact retrieval cannot be tracked to a single locus within the brain. Instead, impairments in arithmetic fact retrieval seem to originate from disconnections of several areas within a left-hemispheric network around the AG, thalamus, STS, STG and MTG and IFG. On a more general level, our study underlines the relevance of future research, taking into account grey matter integrity and white matter connectivity in numerical cognition and cognition in general.

## Acknowledgements

This work was supported by the Deutsche Forschungsgemeinschaft (KL 2788/2-1 und KA 1258/24-1). We thank all patients who participated in this study as well as the nursing staff of the University of Tübingen who assisted with patient preparation.

